# Membrane-active Polymers: NCMNP13-x, NCMNP21-x and NCMNP21b-x for Membrane Protein Structural Biology

**DOI:** 10.1101/2022.01.10.475744

**Authors:** Thi Kim Hoang Trinh, Claudio Catalano, Youzhong Guo

## Abstract

Membrane proteins are a ubiquitous group of bio-macromolecules responsible for many crucial biological processes and serve as drug targets for a wide range of modern drugs. Detergent-free technologies such as styrene-maleic acid lipid particles (SMALP), diisobutylene-maleic acid lipid particles (DIBMALP), and native cell membrane nanoparticles (NCMN) systems have recently emerged as revolutionary alternatives to the traditional detergent-based approaches for membrane protein research. NCMN systems aim to create a membrane-active polymer library suitable for high-resolution structure determination. Herein, we report our design, synthesis, characterization and comparative application analyses of three novel classes of NCMN polymers, NCMNP13-x, NCMNP21-x and NCMNP21b-x. Although each NCMN polymer can solubilize various model membrane proteins and conserve native lipids into NCMN particles, only the NCMNP21b-x series reveals lipid-protein particles with good buffer compatibility and high homogeneity suitable for single-particle cryo-EM analysis. Consequently, the NCMNP21b-x polymers that bring out high-quality NCMN particles are particularly attractive for membrane protein structural biology.

**Graphical abstract:** 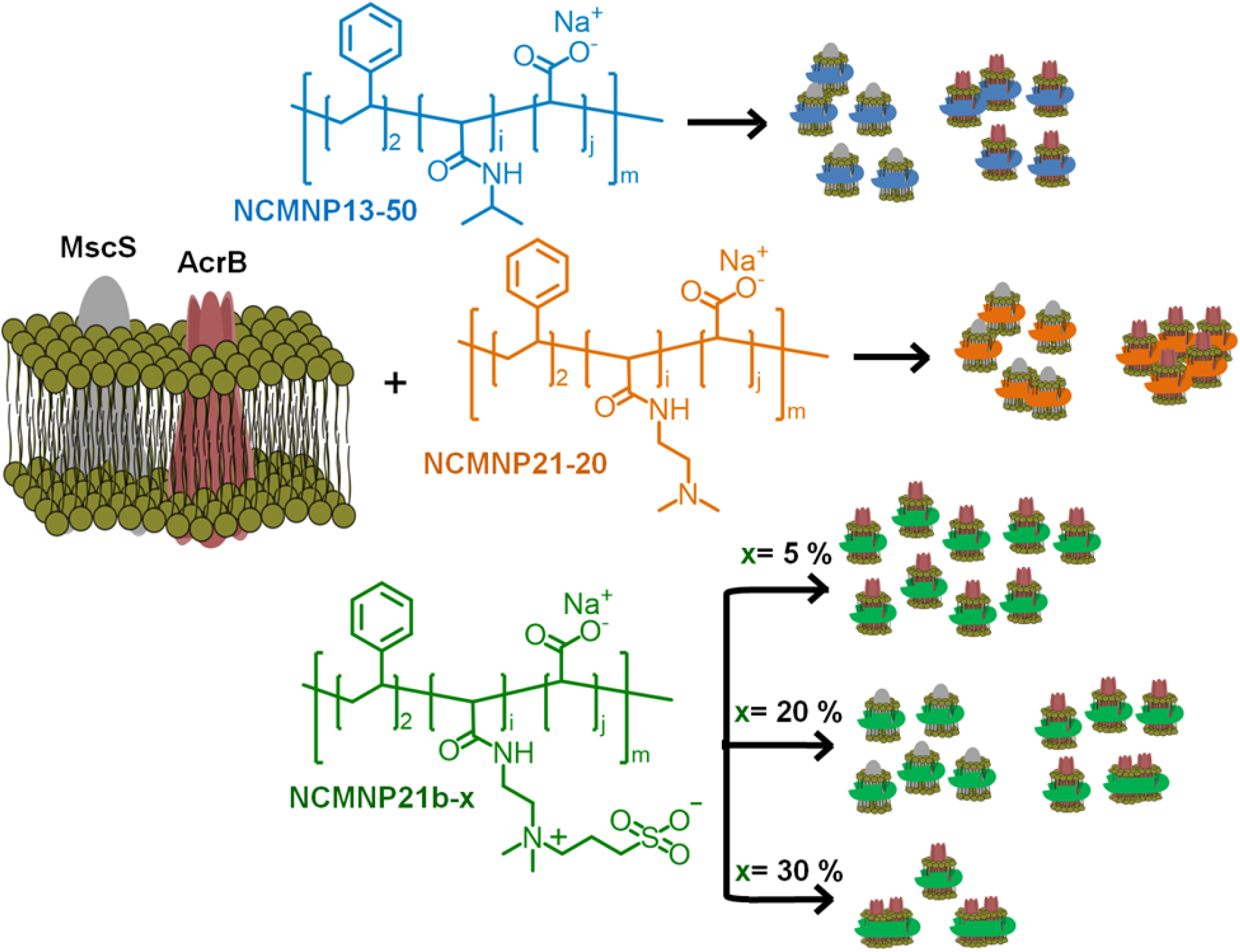

**Highlights:**

- Demonstrate straightforward strategies for tailoring styrene-maleic co-polymer (SMA) that revealed novel buffer compatible polymers, NCMNP13-50, NCMNP21-20 and NCMNP21b-x.
- Elucidate how modification factors alter the membrane-active properties of these polymers, i.e., membrane protein extraction efficiency, morphology, etc.
- Provide valuable insights into the rational design of membrane-active polymers for membrane protein structural biology.
- NCMNP21b-x polymers are highly compatible with high-resolution structure determination using an emerging technique, cryo-EM.

## 1. Introduction

Membrane proteins are an enigmatic group of bio-macromolecules with highly diverse structures and functions involved in many biological processes. Accurate structural information, particularly for membrane proteins involved in protein-lipid interactions, is crucial to understanding active molecular mechanisms.^1^ As leading drug targets, structural and mechanistic comprehension of membrane proteins provides numerous drug development opportunities.^2,3^ Extraction of membrane proteins from the cell membrane environment using detergents is often required for structural analysis. Protein-lipid interactions are often pivotal in maintaining a membrane protein’s natural structure and function. However, detergents often cause over-delipidation during membrane protein solubilization. Synthetic membrane-active polymers can encircle a small portion of the lipid tails without distorting lipid-protein complexes; therefore, they might serve as better alternatives for membrane protein structural studies.^4^ In the polymer-lipid-protein particles, hydrophilic side groups (for example, –COOH) of polymeric belts face toward the aqueous phase that renders the particles suspended and stable for subsequent purification and downstream analysis.^5,6^

Both styrene-maleic acid (SMA) and diisobutylene-maleic acid (DIBMA) co-polymers could preserve and stabilize native membrane protein-lipid complexes.^7–11^ In the case of SMA, upon increasing to a slightly alkaline pH, notable increases in solubilization efficiency of the *Escherichia coli* (*E. coli)* membrane were found. In the basic condition, a lower association among polymeric chains resulting from relatively high negative charge density, in theory, allows them to insert into the lipid bilayer and ultimately induce membrane solubilization.^7,12^ Nonetheless, differences in chemical structure and hydrophobicity of polymeric hydrophobic domains are major determinants influencing yield, size, purity, and long-term stability of resulting particles. Generally, membrane solubilization is mainly driven by the hydrophobic effect.^5^ Thus, it is not surprising when the SMA (2:1) polymer with pronounced hydrophobicity of phenyl groups, as reported by several groups, gave much higher solubilizing efficiency toward some proteins such as ZipA and ABC transporter BmrA than DIBMA, which composes much less hydrophobic portions.^8,13^ DIBMA treatment, in most cases, resulted in protein particles somehow larger in diameter and with less homogeneity.^8,13^ Perhaps the large-size particles accommodated more contaminants and thus were indeed incompatible with structural determination by single-particle cryo-electron microscopy (cryo-EM). Also, SMA particles showed higher stability during long-term storage due to the tight packing of lipid acyl chains related to the insertion and bending of phenyl rings to encompass the native lipid bilayer.^8,13^

With the above superiority, SMA co-polymers have gained much attention in structural studies of membrane proteins. The number of unique membrane protein structures solved increased over 2-fold after the SMA breakthrough and advancements in cryo-EM (2014);^14,15^ however, such a number is still tiny compared with soluble proteins. One of the reasons is that SMA cannot be applied to all membrane proteins, partially because the carboxylate function group could lead to non-covalent aggregation in acidic pH and divalent cation buffers, which are required for some protein families to regulate functions.^12,16^ To overcome current drawbacks, the SMA community has tailored SMA sidechains by using nucleophiles to convert carboxylates into new functional groups such as alcohol (SMA-EA),^17,18^ amino (SMA-ED),^19^ thiol (SMA-SH)^20^ and phosphobetaine (zSMA).^21^ After modification, the stability of particles assembled from these SMA variants, as desired, was partially or entirely turned over; however, none have been demonstrated successful for high-resolution structure determination of membrane proteins with cryo-EM. It is worth to be pointed out that Marconnet et al. have recently revealed a novel cryo-EM compatible polymers, cycloalkane-modified amphipols (CyclAPols), which can provide a native membrane protein extraction but could not retain the well-organized native lipids plug within transmembrane (TM) domains.^22,23^

Among the membrane-active polymers, the commercial SMA2000 with defined ratio 2: 1 of hydrophobic: hydrophilic domains has exhibited as the most effective solubilizer for several membrane protein model systems.^7,9,12,24^ To overcome the limitations of SMA2000 and retain a good extraction efficiency, our novel polymers were produced through individual grafting of isopropylamine (IPA), *N,N*-dimethylethylenediamine (DMEDA), and 3-((2-aminoethyl)-dimethylammonio)propane-1-sulfonate (DMEDA-PS) with various controlled levels (Scheme 1) on SMA2000 side chains. The polymers comprising IPA, DMEDA and DMEDA-PS units were denoted as NCMNP13-x, NCMNP21-x and NCMNP21b-x, respectively, with “x” representing the target grafting level of the amines. While previous studies only employed the grafting level of 50 or 100%, herein, “x” values were varied to explore how such contents and structures of the amines influence membrane solubilization, purification, and stability.

**Scheme 1.**
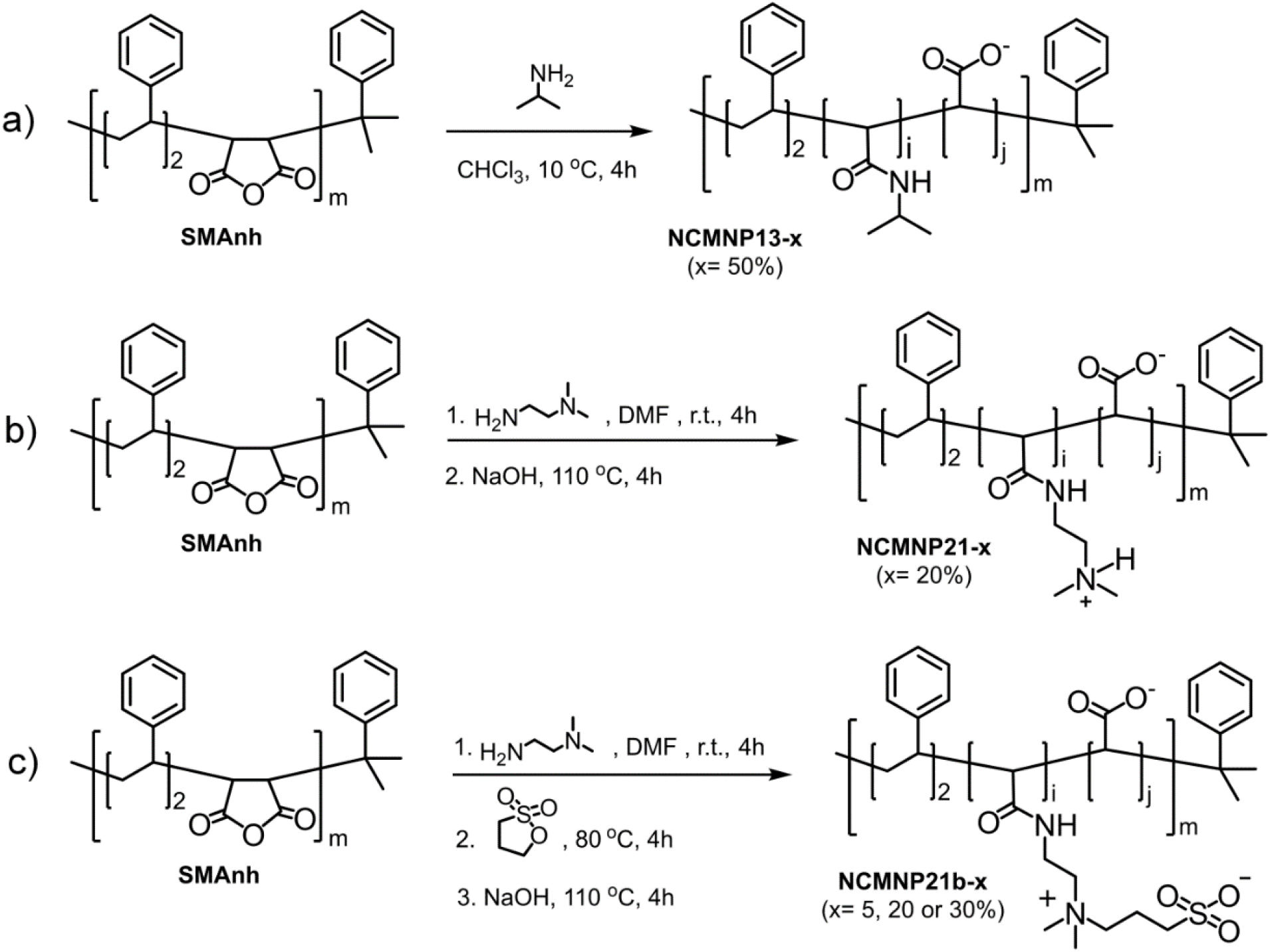
The general synthetic strategy of NCMN polymers. (a) NCMNP13-x. Ispropylamine (IPA) was used with a slight excess amount of maleic anhydride to obtain a complete ring-opening process. (b) NCMNP21-x. (c) NCMNP21b-x.

Here we report our design, synthesis, characterization and comparative application analysis of three new classes of membrane-active polymers: NCMNP13-x, NCMNP21-x and NCMNP21b-x. The comparative application analyses cover four aspects: a) Solubilization efficiency and the long-term stability of NCMN particles. The solubilization efficiency was examined with two different proteins, *E*.*coli* Acriflavine resistance channel protein B (AcrB) and small-conductance mechanosensitive channel protein (MscS). At pH 7.8, AcrB (ExPASy -ProtParam tool calculated pI ∼ 5.35)^25^ has an overall negative charge, and MscS (ExPASy -ProtParam tool calculated pI ∼ 7.9)^25^ has an overall near-neutral charge. That allowed us to elucidate the role of electrostatic interaction for NCMN particle stability. b) Stability of NCMN towards pH conditions and Ca^2+^. c) Lipidomic analysis of NCMN particles. Lipidomic analysis for the AcrB-NCMN particles was performed to explore whether native lipids were encased and possible alterations in their composition. The lipid annulus prevents the collapse of TM domains; thus, confirming their presence is one factor used to justify the suitability of NCMN particles for structural studies. d) Single-particle electron microscopic analysis of NCMN particles. The comparative analyses suggest that NCMNP21b-x has a high potential as a general class of membrane-active polymer for high-resolution structure determination of membrane proteins using single-particle cryo-EM.

## 2. Results and discussions

### 2.1. Design, synthesis and characterization of NCMNP13-x, NCMNP21-x and NCMNP21b-x

The NCMN polymers were prepared via a straightforward ring-opening process of styrene-maleic anhydride (SMAnh) with different nucleophiles and followed by hydrolysis of unreacted maleic anhydrides (MAnhs) with NaOH (Scheme 1). Initially, a fixed molar ratio of amine to MAnh (50% for IPA, 20% for DMEDA) was utilized to prepare NCMNP13-x and NCMNP21-x. For NCMNP21b-x, such a ratio was varied to 5, 20, and 30% to discern their possible effect on the ability to extract membrane proteins. Undergoing hydrolysis afterward made all polymers soluble, which were then precipitated out of the solution to separate from those unreacted residues. The frozen-dried products, as expected, were obtained with high purity, good yields, and were ready to use. The structures of the NCMN polymers were validated using ^1^H NMR and FTIR (Fig. S1– S3).

### 2.2. Comparative application analyses of three class NCMN polymers

#### 2.2.1. Solubilization efficiency and long-term stability of NCMN particles

All membrane solubilization tests were systematically performed with a given set of conditions to perceive modifying factors featuring solubilizing effectiveness and sizes and shapes of achievable NCMN particles.

##### a. Effect of grafted amine nature

Solubilization efficiency was tested using AcrB as a model protein with NCMNP13-50, NCMNP21-20, and NCMNP21b-20, each at a final concentration of 2.5 % (w/v). All membrane suspensions gradually turned clear after the addition of NCMN polymers indicating that these polymers were capable of solubilizing the membrane. Then, the solubilized AcrB and untargeted species were separated using Ni-affinity chromatography (Ni-NTA). All three polymers produced a comparable amount of AcrB-NCMN particles. This is demonstrated by the apparent bands of relatively high similarity in intensity at 110 kDa in SDS-PAGE gels; however, there were other contaminants, despite with much less amount, seen with NCMNP21b-20 (Fig. 1a). Furthermore, whilst the size-exclusion chromatography (SEC) of purified AcrB particles in NCMNP13-50 and NCMNP21-20 were mainly eluted at ∼15 mL, void volume (∼9 mL) was detected in NCMNP21b-20 that implies a larger size of particles (Fig. 1b). The large particles might be responsible for the lower purity in the NCMNP21b-20 sample because the larger portion of lipid assembled can also encapsulate other proteins inside. Yet, it is not a worthy issue for high-resolution structure determination since the impurity could be eliminated by SEC purification, as shown in Fig. S4. These results support the suitability of our functionalized polymers in membrane solubilization and purification.

**Fig. 1.**
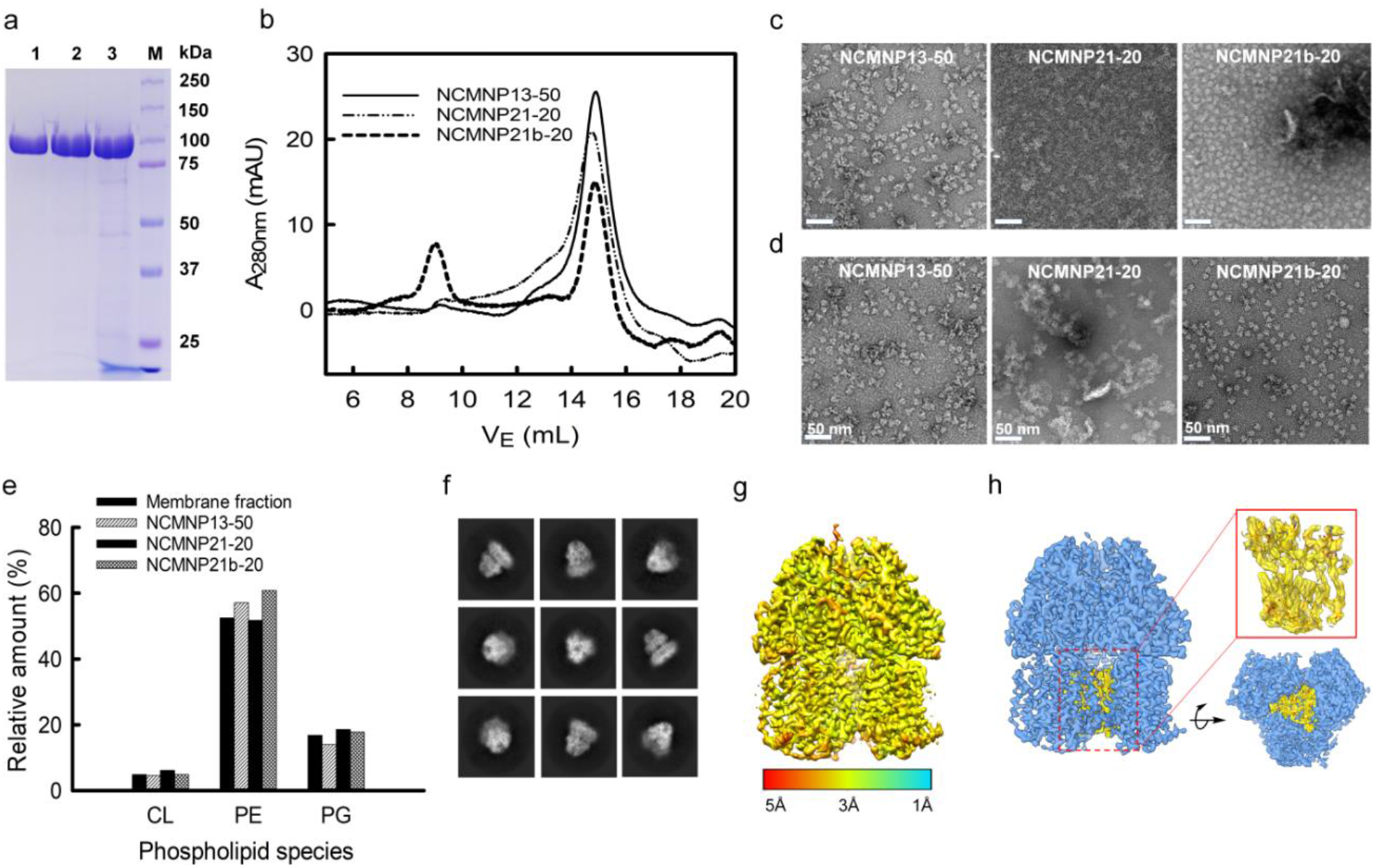
Effect of alkyl amine units of NCMN polymers on the solubilization of AcrB. (a) SDS-PAGE of AcrB particles after Ni-NTA purification visualized by Blue Coomassie (lane 1. AcrB in NCMNP13-50, lane 2. AcrB in NCMNP21-20, lane 3. AcrB in NCMNP21b-20 and M. protein ladder). (b) SEC elution profiles of AcrB particles treated with different NCMN polymers. (c) Negative stain images of one-day-old AcrB particles after Ni-NTA purification (White scale bar represents 50 nm). (d) Negative stain images of one-day-old AcrB particles after SEC purification. (e) Abundance comparison of three major phospholipids in membrane fractions and in AcrB particles displayed as the sum for each of them. (f) Representative 2D class averages of AcrB-NCMNP21b-20. (g) The cryo-EM map of AcrB-NCMNP21b-20 colored by local resolution. (h) Endogenous lipid located with the TM of the pseudosymmetric AcrB trimer.

The size and shape of purified AcrB-NMCN particles were further investigated by negative-stain transmission electron microscopy (TEM) (Fig. 1c–1d). Morphologies of the AcrB-NCMN particles prepared with these polymers are distinctly different. AcrB-NCMNP21-20 particles mainly exist as aggregates, even after the SEC purification. Both AcrB-NCMNP13-50 and AcrB-NCMNP21b-20 particles exist as single particles with a typical triangular shape, with diameters of ∼ 10 nm consistent with the previous particles produced using SMA2000.^9^ As seen, NCMNP21b-20 afforded more monodisperse particle population than NCMNP13-50, making clear the turntable effects of the modified side chains.

The morphological divergence might be caused by undesirable interactions mediated by the chemical nature of the novel head groups, randomly distributed along with polymeric belts, especially in regions unassociated with TM domains. Notably, under our experimental temperature lower than a lower critical solution temperature of NCMNP13-50 (32 °C),^26,27^ it is favored for the nonpolar isopropyl anchors to hydrophobically interact with bound lipids of other neighbor particles that induced the non-covalent aggregation of the NCMN particles.^28^ Similarly, the amino groups of DEADA units incorporated on NCMNP21-20 with a p*K*_a_ value of 8.51 (calculated based on ChemAxon) are probably protonated in the elution buffer pH 7.8, and hence electrostatically interact with carboxylic units, anionic lipids, and cytoplasmic domains,^29^ of which are negative-charged species, consequently destabilizing the NCMN particles. Lastly, in the case of NCMNP21b-20, the less aggregation probably relates to the zwitterionic nature of PS,^30^ where its opposing-charge ions on the same unit associate with each other rather than with other ionic species. By that, the overall charge of the AcrB-NCMNP21b-20 particle surface remains negative, conferring the repulsion between the particles and minimizing their aggregation potential.

#### b. Effect of membrane protein nature

With concerns about whether the efficacy of the NCMN polymers toward the solubilizing of other membrane proteins is the same, we expanded our study to MscS, a member of the mechanosensitive channel family entirely different from AcrB in terms of structure and size. MscS is a homoheptamer, with each monomer having a molar mass of ∼31 kDa and comprising a cytoplasmic domain and a TM domain.^24^ For these three NCMN polymers, there were no significant differences in the yields and degree of purity of MscS-NCMN particles after eluted from the Ni-NTA column as resolved in SDS-PAGE gel (Fig. S5a); however, they gave distinct SEC purification profiles. As shown in Fig. S5b, the elution of AcrB-NCMNP21b-20 emerges as a single peak at ∼14 mL, which is the representative retention volume of MscS-SMA2000.^24^ By contrast, the others show two broad predominant peaks at ∼9 mL and ∼15 mL. The shift to lower elution volume indicates the aggregation of the particles, while the latter is attributed to the disruption of the particle complex. These were supported through further characterization using negative-stain TEM. As shown in Fig. S5c, the NCMNP21b-20 produced uniform particles with high monodistribution, whereas the aggregation was observed for MscS-NCMNP13-50 and MscS-NCMNP21-20 samples at different levels. More aggregation was found for NCMNP21-20, which, as explained in the case of AcrB-NCMN particles, is because of the electrostatic interactions between positively charged side groups of DMEDA units and negatively charged clusters located in other MscS-NCMNP21-20 particles nearby. Such electrostatic interactions are much stronger than in between MscS-NCMNP13-50 particles, with only non-specific hydrophobic interactions occurring at IPA segments. Surprisingly, the MscS complexes purified by NCMNP13-50 and NCMNP21-20 might be broken during the SEC purification, evidenced by smaller undefined-shaped particles on the TEM micrograph (Fig. S5c–S5d). The flexibility of MscS-TM domains might cause a reduction in binding affinity to the polymers. The loose binding between the protein particles and polymers might be further damaged when the NCMN particles go through the SEC column because of mechanical shearing forces. In contrast, regardless of aggregation, the well-defined shape of MscS can be seen in the SEC elution fraction of NCMNP21-20 (Fig. S5d). We surmise that at pH 7.8, the negative charge on MscS with the isoelectric point (pI) of ∼ 7.9 is less than that on AcrB (pI ∼ 5.35). Of course, that could also be because of the weak electrostatic interaction with amino groups on NCMNP21-20, thus lowering particle aggregation. These results suggest that the solubilizing behavior of NCMN polymers may be protein-independent, but the long-term stability of the resulting NCMN particles remarkably relies on the nature of both the NCMN polymers and membrane proteins.^29^

#### c. The effect of grafting percentage

NCMNP21b-20 produced better NCMN particles than NCMNP13-50 and NCMNP20-20; therefore, NCMNP21b-5 and NCMNP21b-30 were used to examine the influence of the grafting degree on the protein solubilization. The solubilization results tell us that grafting percentage dramatically changes the purification yield, purity and particle morphology (Fig. 2). Fascinatingly, the highest yield and purity of AcrB particles were attained with NCMNP21b-5 and decreased in the following order: NCMNP21b-5 > NCMNP21b-20 > NCMNP21b-30. Like SMA2000, NCMNP21b-5 produced only single particles with roughly 10 nm diameters. The major of AcrB-NCMNP21b-20 were small single particles, while the larger patches were prevailed for AcrB-NCMNP21b-30. It seems that the ability to assemble into single particles decreases as the grafting levels of PS increases. However, it begs the question as to why the NCMNP13-50 contains 50% of the IPA units but gave solely single particles. Back to the work of Scheidelaar et al. depicting the SMA-induced solubilization mechanism,^5^ the efficient action of SMA is mainly driven by hydrophobic interactions and satisfied with a relatively small cross-sectional area given by its –COOH side groups. Fundamentally, the hydrophobic insertion of the phenyl ring into the lipid bilayer, known as a crucial prerequisite for membrane assembly, is favorable with –COOH as a neighbor unit because it can minimize the steric hindrance. Replacement of these typical units by larger alkyl groups probably creates a relatively large cross-sectional area that can impede the phenyl-lipid interactions; consequently, resulting in the larger size and lower yield of particles.^31–34^ As described here, despite having a high number of –COOH replaced, NCMNP13-50 still could give the single particles since its IPA units are small enough not to give notable negative effects. Otherwise, the high grafting degree of PS, a much larger group than –COOH and IPA, made NCMNP21b-x challenging to insert into the membrane to trigger the solubilization, bringing out a decline in the solubilization efficacy. These findings help us understand the correlation between modification levels and membrane solubilization action, hence paving a way to tailor NCMN polymers to reach the goals of future studies optimally.

**Fig. 2.**
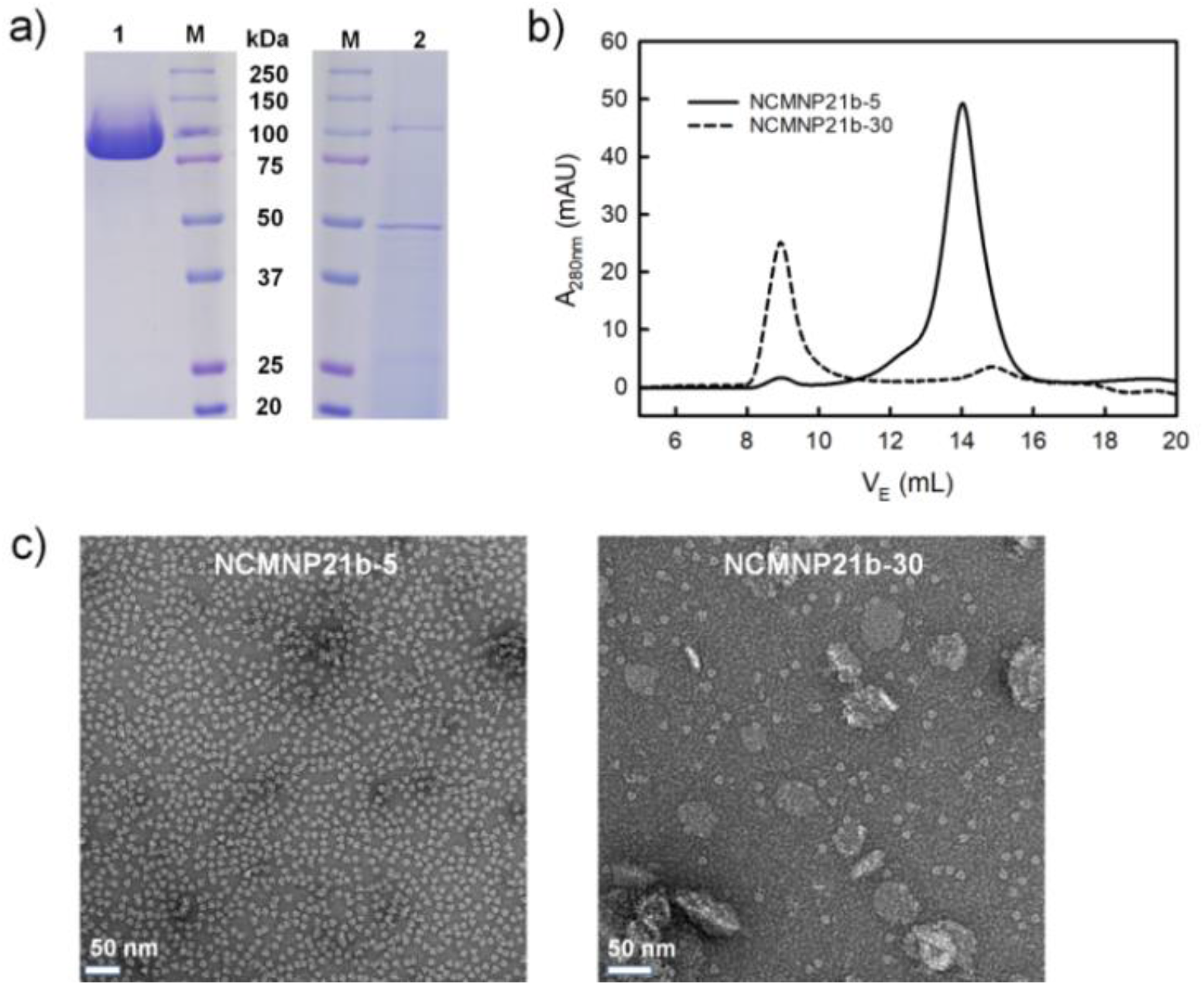
Effect of the grafting degree of NCMN polymers on the solubilization of AcrB. (a) SDS-PAGE of AcrB particles after Ni-NTA purification visualized by Blue Coomassie (lane 1. AcrB in NCMNP21b-5, lane 2. AcrB in NCMNP21b-30 and M. protein ladder). (b) SEC elution profiles of AcrB particles treated with different NCMN polymers. (c) Negative stain images of one-day-old AcrB particles after Ni-NTA purification (White scale bar represents 50 nm).

#### 2.2.2. Stability of NCMN particles toward pH and Ca^2+^

Next, we measured the turbidity of the AcrB-NCMN particles to assess how our modification is effective for improving pH and divalent cation compatibility (Fig. S6–S7). On account of the short-term stability of some particles, all measurements were performed immediately after the Ni-NTA purification. Current instability problems of SMA, which, as apprehended, relate to the presence of –COOH groups. Therefore, similar to prior studies,^19,21,35^ substituting –COOH units by the selected amines showed the capabilities of turning the particle stability as summarized in Table 1.

**Table 1.** Structure composition of the customized NCMN polymers and the stability of its AcrB particles after membrane solubilization.

| Polymer | Amine/ MAnh<br>feeding ratio<sup>a</sup> | Grafting degree<br>of amine (%)<sup>b</sup> | Optimum<br>pH<sup>c</sup> | Tolerance to [Ca<sup>2+</sup>]<br>(mM)<sup>c</sup> |
| --- | --- | --- | --- | --- |
| NCMNP13-50 | 1.05 | 47.4 | > 3 | ≤ 7.5 |
| NCMNP21-20 | 0.40 | 17.7 | > 3 | ≤ 2.5 |
| NCMNP21b-5 | 0.10 | 4.4 | > 4 | ≤ 5.0 |
| NCMNP21b-20 | 0.40 | 18.0 | > 3 | ≤ 15.0 |
| NCMNP21b-30 | 0.60 | 27.5 | > 2 | ≤ 25.0 |

<sup>a</sup>Molar ratio. <sup>b</sup>Experimental grafting percentage of amine calculated based on <sup>1</sup>H NMR characterization.
<sup>c</sup>Stability of AcrB particles in various buffer environments determined based on turbidity experiments (Fig. S6 – S7).

The AcrB-NCMNP13-50 particles begin to aggregate at pH below 3 and in the presence of [Ca^2+^] higher than 7.5 mM, indicated by the increased absorption bands (Fig. S6a, Fig. S7a). The compatibility of wider pH ranges and divalent cations compared to SMA2000 is most likely due to approximately 50% of the negative charge reduced on the NCMNP13-50 polymer.^6^ Meanwhile, albeit having lower Ca^2+^-tolerance capability than NCMNP13-50, the NCMNP21-20 particles can resist pH down to pH 3 even with only 20% of DMEDA supplants –COOH (Fig. S6b, Fig. S7b). This enhanced acidic pH stability might be attributed to the presence of positively charged groups, which can somehow preclude non-covalent aggregation resulting from protonation of –COOH groups.^19,35,36^ Taking a deep look into the NCMNP21b-x series revealed that with the same grafting levels as NCMNP21-20, the NCMNP21b-20 particles exhibit better stability toward Ca^2+^ with a concentration up to 15 mM. This active concentration is lower for NCMNP21b-5 (5 mM) and higher for NCMNP21b-30 (25 mM) (Fig. S7c–S7e). A similar trend is observed in the pH tolerance, which increases along with the grafting degree (Fig. S6c–S6e). The presence of pH-insensitive PS units probably leads to the formation of negative-charge holes on the belt surface, allowing Ca^2+^ to be complexed rather than its affinity binding with –COOH groups.^37–39^ Therefore, it could account for the notable improvement in Ca^2+^ resistance of NCMNP21b-x. In conclusion, the stability of NCMN particles is, in fact, dependent on the nature of the side groups and the grafting percentage.

### 2.2.3. Lipidomic analysis of NCMN particles

Even though 24 lipid molecules were found encased in AcrB-SMA2000,^9^ whether or not preferential lipid-polymer bindings exist remains unclear. Herein, we shed light on the lipid composition to explore the possible effect of polymeric sidechains on lipid selectivity through mass spectrometry (MS).^40^ As shown in Fig. 1e, phosphatidylethanolamine (PE), phosphatidylglycerol (PG), and cardiolipin (CL) are the major lipid species. The lipids identified are comparable in extracted samples and consistent with earlier reports of the lipids in the *E*.*coli* membrane.^41^ The results suggest that NCMN polymers preserve the native lipids in NCMN particles and indicate that none of the polymers interacts with polar head groups of the lipids. Nevertheless, apparent variability in length and saturation of associated acyl chains are observed. The lipid molecules with long carbon tail and high degree of unsaturation become more abundance in the purified AcrB particles (Fig. S8). Since this enrichment is found in all the NCMN particles with relatively high similarity, it suggests that AcrB intrinsically has preferred association with those long chain and unsaturated natural lipids, and that variation of polymeric side chains on the NCMN polymers do not perturb the protein-lipid interactions.

### 2.2.4. Single-particle electron microscopic analysis

Comparative analysis of the AcrB-NCMN particles on the negative-stain TEM micrographs clearly shows that NCMNP21b-x polymers can bring out much more homogenous NCMN particles at the grafting percentages between 5% and 20%. Consequently, such uniform particles were further analyzed by single-particle cryo-EM. Fig. 1f displayed 2D class averages of AcrB-NCMNP21b-20 with high similarity to AcrB-SMA2000.^9^ Following 3D classification, 3D refinement with employed C1 asymmetry revealed a final EM density map at a 3.69 Å global resolution (Fig. 1g and Fig. S9). That allowed for improved clarification of the secondary structure of AcrB protein and co-purified lipids (Fig. 1h) residing in the center cavity of cytoplasmic domains, distinctly. Our results suggest that NCMNP21b-20 polymer is superior to SMA2000 in wide range pH a divalent cation ion compatibility and more advanced than CyclAPol in retaining native cell membrane lipids during membrane solubilization and protein purification.^9,23^

## 3. Conclusions

In this contribution, we described novel SMA variants referred to as NCMN polymers and methodically investigated the effects of their sidechains on the membrane solubilization, morphology, and stability of protein particles. These polymers were synthesized by substituting – COOH groups of commercial SMA2000 with various alkyl amide units. Systematical comparison of these polymers in application to two different membrane proteins revealed that:

- All NCMN polymers can directly trigger the solubilization of AcrB and MscS proteins, which differ in size and structure, into NCMNs.
- The coupling levels of amines mainly drive the size and homogeneity of the AcrB-NCMN particles. A high number of large alkyl side groups can interfere with styrene-lipid interaction that lowers the chance to give the high yield of small single particles.
- The long-term stability of NCMN particles depends on the chemical nature of polymers and protein structure. In principle, with much less negative charge on the surface at pH 7.8, MscS-NCMN particles resulted from NCMNP21-20 bearing a positive charge on a sidechain that exhibited more advanced stability AcrB-NCMN particles in the same polymer.
- Coupled with more non-chelating units, the NCMN polymers become more compatible with broad range pH conditions and divalent cations.
- Side groups do not interact with lipids enabling the NCMN polymers to retain the lipids in NCMN particles with their composition as close as in the native state.
- NCMNP21b-x polymers bearing much better divalent ion and broader pH range compatibility are particularly attractive for membrane protein structural biology.

These findings provide valuable insights into crucial factors that allow us to purposefully engineer SMA to provide desirable properties for structural biology studies in the future.

## 4. Methods

### 4.1. Materials

Styrene-maleic anhydride copolymer (SMAnh, St: MAnh= 2:1, *M*_n_ = 3,000 g·mol^-1^, *M*_w_ = 7,500 g·mol^-1^, acid number: 358 g KOH·mol^-1^) was purchased from Cray Valley USA, LLC (Exton, PA). *N,N*-Dimethylformamide (DMF, 99.8%, extra dry, anhydrous, AcroSeal), chloroform (CHCl_3_, 99.9%, extra dry, stabilized, AcroSeal) and deuterium oxide (D_2_O, 99.8 atom % D) were brought from Acros. Isopropylamine (IPA, > 99%, reagent grade), 1,3-propanesultone (PS, 98%) and tris (2-carboxyethyl) phosphine hydrochloride (TCEP) were obtained from TCI, Sigma and GoldBio, respectively. Sodium hydroxide (NaOH, pellets, certified ACS), calcium chloride dihydrate (CaCl_2_.2H_2_O, certified ACS), sodium chloride (NaCl, crystalline, certified ACS), diethyl ether (laboratory), hydrochloric acid (HCl, 36.5 to 38.0%, certified ACS plus), sodium acetate anhydrous (white crystals), glycine (white crystals), HEPES (white crystals, molecular biology) and imidazole (molecular biology), were bought from Fisher. All chemicals were used as received.

### 4.2. Polymer synthesis

#### NCMNP13-50 synthesis

IPA (1,373 µL, 16.78 mmol, 1.05 equiv.) in 10 mL of CHCl_3_ was added dropwise into a vigorously stirred solution of SMAnh (5 g, 15.85 mmol of MAnh, 1 equiv.) in 20 mL of CHCl_3_ at 0 °C. The mixture was reacted at room temperature for 4 h, followed by removing the solvent. The collected powder was dissolved in NaOH (0.6 M) and then precipitated with HCl (12 M). The polymer was then pelleted through a centrifuge at 12,000 × *g* for 20 min. Subsequently, the HCl was decanted, and the polymer was rinsed again with water (2 times). Finally, the polymer was re-dissolved in NaOH (0.6 M), and the pH of the polymeric solution was adjusted to 7.8 before lyophilization.

#### NCMNP21-20 synthesis

DMEDA (692 µL, 6.34 mmol, 0.4 equiv.) in 10 mL of DMF was added dropwise into a vigorously stirred solution of SMAnh (5 g, 15.85 mmol of MAnh, 1 equiv.) in 20 mL of DMF at room temperature. The mixture was then reacted at room temperature for 4 h followed by the precipitation of the polymer with excess diethyl ether. After being dried at 25 °C in a vacuum oven, the white powder was mixed with 25 mL of NaOH (1 M) and heated to reflux for 4 h. The transparent solution was cooled down to room temperature before being precipitated with HCl (12 M). Subsequently, the polymer was pelleted through a centrifuge at 12,000 × *g* for 20 min. Afterward, the HCl was decanted, and the polymer was rinsed again with water (2 times). Finally, the polymer was re-dissolved in NaOH (0.6 M), and the pH of the polymer solution was adjusted to 7.8 before lyophilization.

#### NCMNP21b-20 synthesis

DMEDA (6.92 µL, 6.34 mmol, 0.4 equiv.) in 10 mL of DMF was added dropwise into a vigorously stirred solution of SMAnh (5 g, 15.85 mmol of MAnh, 1 equiv.) in 20 mL of DMF at room temperature. The mixture was reacted at room temperature for 4 h prior to PS (584 µL, 6.66 mmol, 0.42 equiv.) addition. The solution was then placed in an 80 °C oil bath and stirred for 4 h. The obtained solution was precipitated with excess diethyl ether and dried at 25 °C in a vacuum oven. The powder was mixed in 25 mL of NaOH (1 M) and heated to reflux for 4 h. After cooling to room temperature, the transparent solution was precipitated with HCl (12 M). The polymer was pelleted through a centrifuge at 12,000 × *g* for 20 min. Afterward, the HCl was decanted, and the polymer was rinsed again with water (2 times). Finally, the polymer was re-dissolved in NaOH (0.6 M), and the pH of the polymer solution was adjusted to 7.8 before lyophilization.

#### NCMNP21b-5 and NCMNP21b-30 synthesis

These polymers were prepared following the NCMNP21b-20 protocol. Nevertheless, the amount of DMEDA and PS were altered as mentioned below:

– NCMNP21b-5: DMEDA (172 µL, 1.58 mmol, 0.1 equiv.) and PS (306 µL, 3.49 mmol, 0.22 equiv.)
– NCMNP21b-30: DMEDA (1,038 µL, 9.52 mmol, 0.6 equiv.) and PS (862 µL, 9.84 mmol, 0.62 equiv.)

### 4.3. Polymer characterization methods and instruments

#### Fourier-transform infrared (FTIR) spectroscopy

The FT-IR spectra were recorded on a Nicolet iS10 spectrometer (Thermo Scientific) equipped with a smart diamond ATR accessory. All absorbance spectra were gathered with 32 scans with a resolution of 4 cm^-1^ in the range of 4000 – 650 cm^−1^.

#### Nuclear magnetic resonance (NMR) spectroscopy

All NMR experiments were performed on a Bruker Fourier 300 spectrometer with a frequency of 300 MHz. Chemical shifts (δ) are declared in ppm in relation to the solvent residual peak of D_2_ O at 4.79 ppm.

#### Ultraviolet-visible (UV-vis) spectroscopy

UV-vis spectra were recorded on Micro-Volume Measurement spectrometer MMC-1600C obtained from Shimadzu. The measurements were conducted using 200 µL of the sample with a scanning speed of 400 nm·min^-1^.

### 4.4. Expression and purification of AcrB and MscS

Both 8 His-tagged AcrB and 10 His-tagged MscS were over-expressed in the outer membrane of the *E*.*coli*, and their membrane fractions were prepared as described previously.^24,42,43^ Following a general solubilization protocol, 1 g membrane fraction was suspended and homogenized in 10 mL NCMN Buffer A using a Dounce homogenizer. Afterward, the homogenized membrane was transferred to a 50 mL polypropylene tube and mixed with NCMN polymer for a final concentration of 2.5% w/v. The incubation was carried out for 2 h at 20 °C, and the insoluble species were then spun down at 200,000 × *g* for 1 h at 20 °C. The supernatant was collected and loaded onto a 5 mL Ni-NTA column (GE Healthcare) pre-equilibrated with NCMN Buffer A at a flow rate of 0.5 ml·ml^-1^. The column was washed with 25 mL of each NCMN Buffer B and NCMN Buffer C prior to the elution of the protein with 20 mL of a mixture buffer of the NCMN Buffer C and NCMN Buffer D (1:1 v/v). Collected fractions containing membrane proteins were loaded onto a Superose 6 increase 10/300 column (GE Healthcare) and eluted with 30 mL of NCMN Buffer E. The NCMN polymer was added to all washing and elution buffers with a final concentration of 0.05% (w/v).

All the buffers were filtered with 0.22 µm MCE Membrane (MF-Millipore™) before using, and their compositions are listed below:

– NCMN Buffer A: 50 mM HEPES, pH 7.8, 500 mM NaCl, 5% glycerol, 20 mM imidazole, 0.1 mM TCEP
– NCMN Buffer B: 25 mM HEPES, pH 7.8, 500 mM NaCl, 40 mM imidazole, 0.1 mM TCEP
– NCMN Buffer C: 25 mM HEPES, pH 7.8, 500 mM NaCl, 75 mM imidazole, 0.1 mM TCEP
– NCMN Buffer D: 25 mM HEPES, pH 7.8, 500 mM NaCl, 500 mM imidazole, 0.1 mM TCEP
– NCMN Buffer E: 40 mM HEPES, pH 7.8, 500 mM NaCl, 0.1 mM TCEP.

### 4.5. NCMN particles characterization methods and instruments

#### Stability of NCMNs toward pH-and divalent cation-based buffers

To testify the AcrB-NCMN particle stability, different buffers were prepared with the composition listed below:

– Ca^2+^-based buffers (pH 7.8): 40 mM HEPES, 100 mM NaCl, 0.1 mM TCEP and a given CaCl_2_ concentration (0 – 100 mM).
– pH buffers:
– Tris buffer (pH 10): 40 mM Tris, 100 mM NaCl, 0.1 mM TCEP
– HEPES buffer (pH 7.8): 40 mM HEPES, 100 mM NaCl, 0.1 mM TCEP
– Sodium acetate buffer (pH 4 and 5): 40 mM sodium acetate, 100 mM NaCl, 0.1 mM TCEP
– Glycine buffer (pH 2 and 3): 40 mM glycine, 100 mM NaCl, 0.1 mM TCEP.

Ca^2+^ sensitivity. To 100 µL of AcrB-NCMN particles (OD_280_ ∼ 1 mg/ mL) in elution buffer (pH 7.8) was added dropwise 100 µL of Ca^2+^-based buffer to achieve the final Ca^2+^ concentration (0 – 50 mM). After 1 h incubation, the transparent changes were monitored by a NanoDrop spectrophotometer.

pH tolerance. To 100 µL of AcrB-NCMN particles (OD_280_ ∼ 1 mg/ mL) in elution buffer (pH 7.8) was added dropwise HCl 1M (or NaOH 1M) to alter its pH to desirable pH, followed by the addition of corresponding pH buffers to reach a final volume of 200 µL. The mixtures were incubated for a further 1 h. Finally, the transparent changes were monitored by a NanoDrop spectrophotometer.

All experiments were carried out in a NanoDrop™2000 spectrophotometer (Thermo Scientific). All spectra with a normalized 10 mm pathlength absorbance were collected in the preconfigured Protein A280 method with a bandwidth of 1 nm and a scan speed of 800 nm·min^-1^.

#### Transmission electron microscopy (TEM)

Briefly, a 3.5 µL of the sample (OD_280_ ∼ 0.1 mg/mL) was absorbed to a 400 mesh carbon-coated copper grid, which was glow-discharged at 20 mA for 1 min. After washing 3 times with water, and twice with 2% (w/v) uranyl acetate, the sample was stained with 2% (w/v) uranyl acetate for 1 min. Removal of the excess stain was carried out by touching the edge of the grid to a piece of Whatman filter paper. After air drying, the grid was imaged using a transmission electron microscope (Tecnai F20, UVA) at 62,000 × magnification at specimen level.

#### Lipid analysis

The endogenous lipids were extracted as described previously with the following modifications.^44^ 1 volume of AcrB-NCMN particles (OD_280_ ∼ 1 mg/ mL) was mixed with 2 volumes of chloroform: methanol (2:1 v/v) and allowed to shake 1 h at 4 °C. Afterward, the mixture was centrifuged at 12,000 × *g* and 4 °C for 10 min to give phase separation. The aqueous layer (upper phase) was removed, while the organic layer (lower phase) was washed twice with 1 volume of cold water and evaporated to complete dryness. The total lipid extract was analyzed by the Thermo Scientific™ Vanquish™ UHPLC system utilizing a C18+ 2.1 (i.d.) x 150 mm reverse-phase column with 1.5 µm particles. The mass spectrometer was operated at 55 °C with a binary solvents system, mobile phase A1 (CH_3_CN/H_2_O, 50/50, v/v, with 5 mM ammonium formate and 0.1% formic acid) and mobile phase B1 (CH_3_CHOHCH_3_/CH_3_CN/H_2_O, 88/10/2, v/v/v, with 5 mM ammonium formate and 0.1% formic acid) at a flow rate of 0.26 mL/min. The lipid sample was characterized in both positive and negative ionization modes. Full mass spectra were recorded in a range of 300 to 2000 m/z with the resolution set to 30,000. All data were analyzed via Thermo Scientific’s Lipid Search 4.2 software.

#### Single-particles cryo-EM grid preparation, data collection, processing, and 3D EM map reconstruction

Samples were prepared for cryo-EM by applying 3.5 µL of freshly purified AcrB NCMNP21b-20 to glow-discharged Quantifoil R 1.2/13, 300 mesh gold grids. The sample was blotted for 25 s with a force of 6 and then flash-frozen in liquid ethane and stored in liquid nitrogen using a Vitrobot Mark IV (Thermo Fisher Scientific, Waltham, USA) with the environmental chamber set at 100% humidity, 4 °C. Cryo-EM specimen grids were imaged on a Titan Krios operated at 300 kV equipped with a Gatan K3 direct electron detector camera at New York Structural Biology Center. Images were taken at 105,000 × nominal magnification, corresponding to a calibrated pixel size of 0.8256 Å /pixel. An initial dataset of 8,591 micrographs for AcrB was obtained by automated data collection using Leginon, with nominal defocus values ranging between 0.4 and 2.9 µm at a dose rate of 44.18 e-/Å 2/s with a total exposure of 1.36 s, for an accumulated dose of 60.22 e-/Å 2.^45^ The full images dataset was processed using CryoSparc v3.3.1: beam-induced motion and CTF were corrected using patch motion and CTF correction function modules.^46,47^ The exposures were manually curated, removing all the micrographs with more than 1,000 Å astigmatism and with more than 5 Å CTF fit resolution. ∼1.5 millions particles were automatically picked with Topaz and extracted with a box size of 360.^48^ Several rounds of 2D classifications were performed to remove junk particles and duplicates. The best-looking 2D classes and their respective particles were then subjected to ab-initio reconstruction. Non-uniform reconstruction was then performed on 85,980 particles with C1 symmetry, and the resulting sharpened model had a resolution of 3.6 Å according to the gold-standard Fourier shell correlation. (Fig. S9). The Cryo-EM data collection and processing parameters are summarized in Table S1.

## Author Contributions

YG conceived the project. YG and TKHT designed the experiments. TKHT performed the polymer synthesis, characterization, NCMN sample preparation, analysis, and cryo-EM data processing. C.C participated in growing cell culture and preparation of the cryo-EM map. YG oversaw and supervised the project. TKHT drafted the manuscript. YG, TKHT and CC revised the manuscript.

## Acknowledgements

YG was supported by the VCU School of Pharmacy and Department of Medicinal Chemistry through startup funds, the Institute for Structural Biology, Drug Discovery and Development through laboratory space and facilities, and NIH Grant R01-GM132329 (To YG). The funders had no role in study design, data collection, analysis, decision to publish, or manuscript preparation. The content is solely the authors’ responsibility and does not necessarily represent the National Institutes of Health or other funding organizations’ official views. The lipidomic analyses were performed at the VCU Lipidomics/Metabolomics Core with the NIH-NCI Cancer Center Support Grant P30 CA016059 to the VCU Massey Cancer Center and shared resource grant (S10RR031535) from the National Institutes of Health. We also thank ChemAxon for providing us with the academic license to access ChemAxon 20.11.0 version for p*K*a calculation. We are grateful to Edward T. Eng, Charlie Dubbeldam, Carolina Hernandez, Kashyap Maruthi for their great help for cryo-EM data collection at the National Center for CryoEM Access and Training (NCCAT) and the Simons Electron Microscopy Center located at the New York Structural Biology Center, supported by the NIH Common Fund Transformative High-Resolution Cryo-Electron Microscopy program (U24 GM129539), and by grants from the Simons Foundation (SF349247) and NY State Assembly.

## Supplementary materials

Supplementary materials associated with this article can be found in the online version.

