## Supporting information for "Membrane-active Polymers: NCMNP13-x, NCMNP21-x and NCMNP21b-x for Membrane Protein Structural Biology"

### I. Additional figures

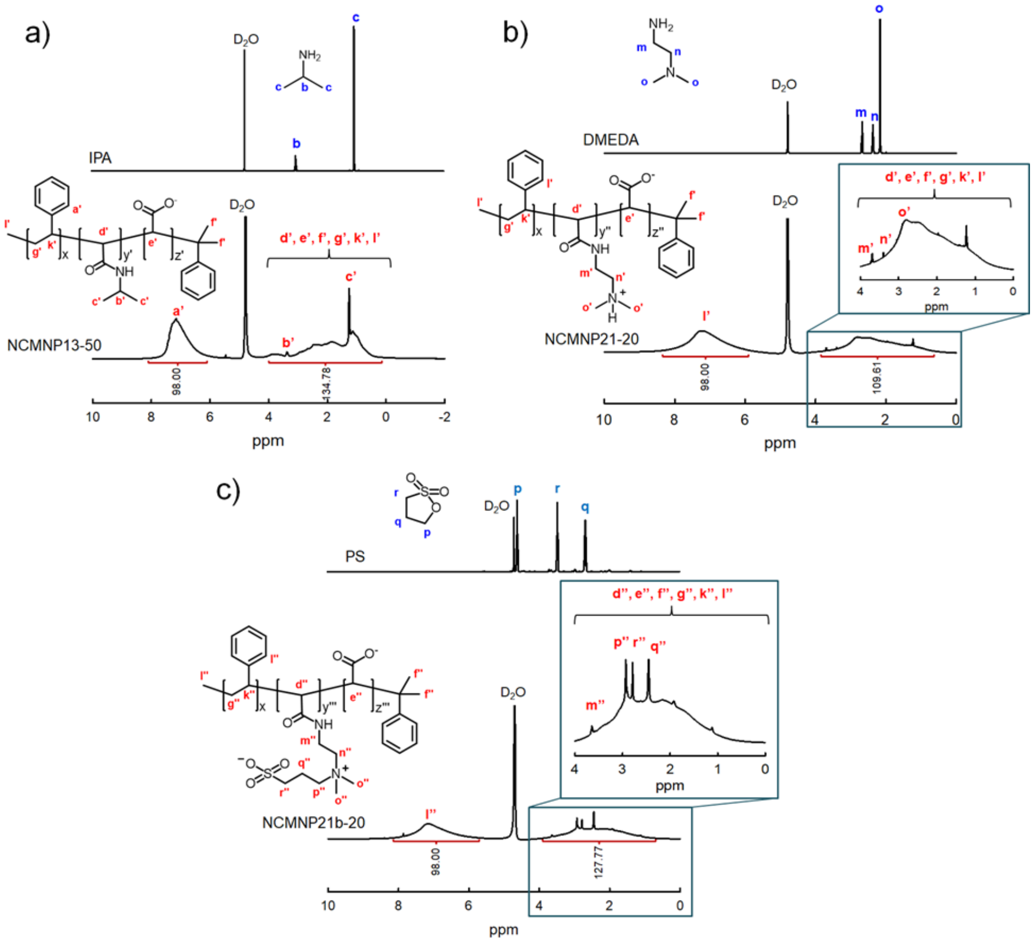

**Figure S1.** $^1$H NMR spectra of NCMN polymers and their modification reactants with their characteristic signals in line with previous studies.[1–3] (a) IPA and NCMNP13-50. The coupling reaction led to a complete shift of the methine and methyl protons from 3.04 ppm and 1.06 ppm (in IPA) to 3.38 ppm and 1.27 ppm (in NCMNP13-50). (b) DMEDA and NCMNP21-20. The downfield shifts of methylene ($H_{m'}$ and $H_{n'}$) and methyl ($H_{o'}$) protons of NCMNP21-20 compared to its in DMEDA were observed, indicating the formation of the target structure. (c) PS and NCMNP21b-20. Proton $H_p$ at 4.69 ppm of PS entirely disappeared in the NCMNP21b-20 spectrum. Meanwhile, new signals assigned to the methylene protons ($H_{p''}$, $H_{q''}$ and $H_{r''}$) of the sulfobetaine fragments were detected with the correct number of aliphatic protons expected for the customized NCMNP21b-20, confirming success in gaining the precise control of grafting degree.

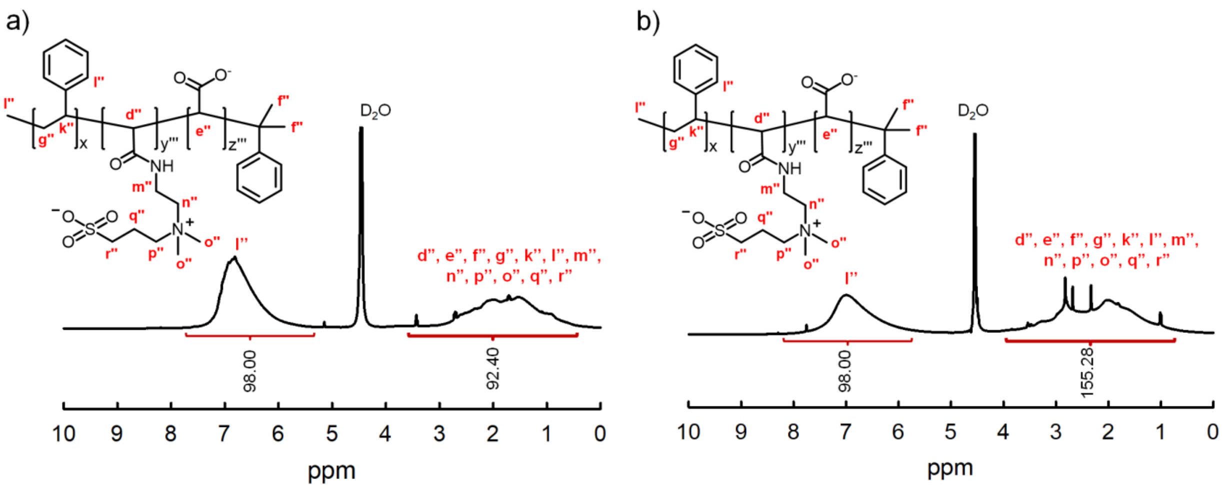

**Figure S2.** $^{1}$H NMR characterization of NCMNP21b-x with full integration. (a) NCMNP21b-5. (b) NCMNP21b-30.

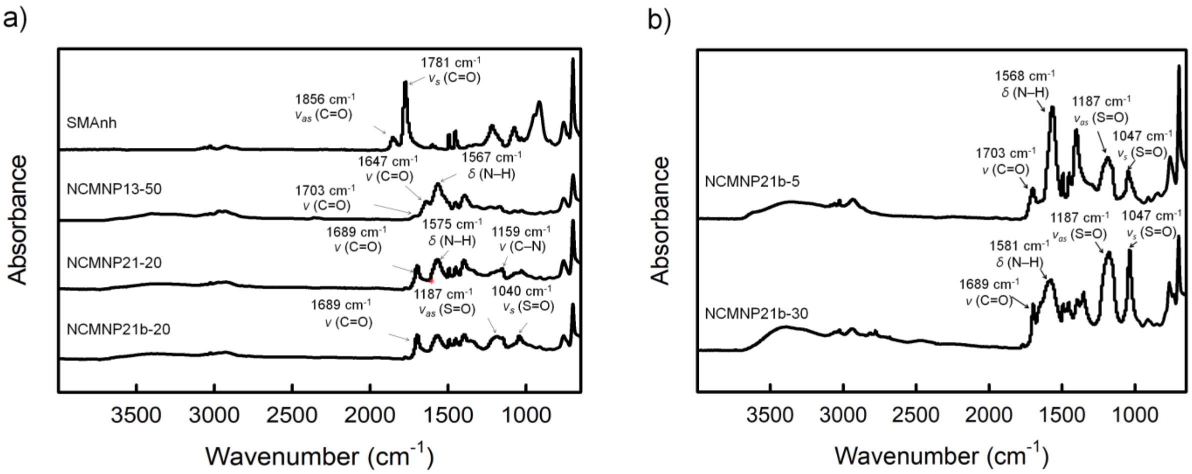

**Figure S3.** FTIR characterization with ATR mode. (a). Spectra of SMAnh, NCMNP13-50, NCMNP21-20 and NCMNP21b-20. All characteristic bands marked are congruent with previous works.[4–7] The red asterisk illustrates a broad overlapping signal of amide N–H bending and sulfobetaine C–N$^{+}$ stretching (1575 cm$^{-1}$). Its center varies depending on the grafting percentage of PS (Figure S2). (b) Spectra of NCMNP21b-5 and NCMNP21b-30. An increase in vibration signals of sulfonate S=O and shift of overlapping peaks of quaternary amine C-N$^{+}$ and amide N-H bending were sought in NCMNP21b-30. It suggests the higher amount of PS was grafted on the backbone of NCMNP21b-30.

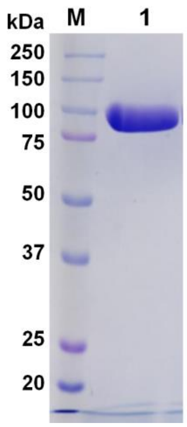

**Figure S4**. SDS-PAGE gel of AcrB in NCMNP21b-20 after SEC purification.

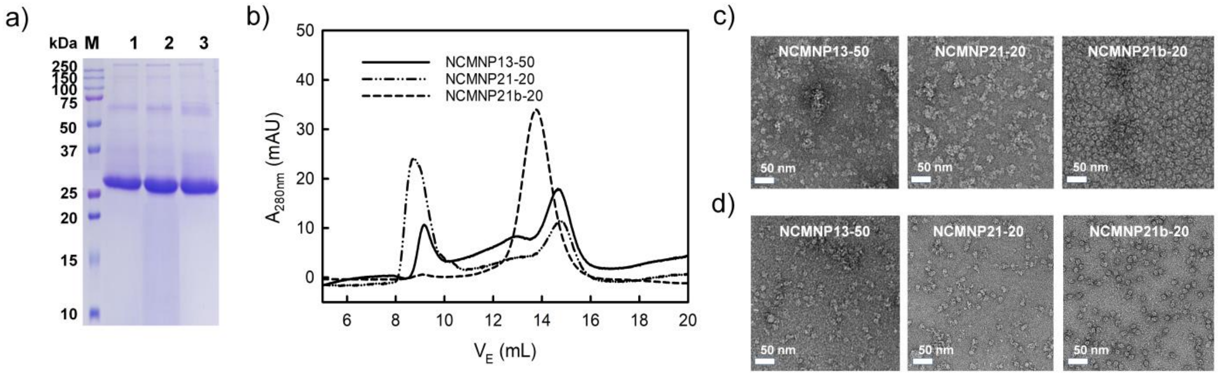

**Figure S5**. Solubilization of MscS protein by various NCMN polymers. (a) SDS-PAGE of MscS particles after Ni-NTA affinity column purification visualized by Blue Coomassie (M. protein ladder, lane 1. MscS in NCMNP13-50, lane 2. MscS in NCMNP21-20 and lane 3. MscS in NCMNP21b-20). (b) SEC elution profiles of MscS particles wrapped by different NCMN polymers. (c) Negative stain images of one-day-old MscS particles after Ni-NTA purification (White scale bar represents 50 nm). (d) Negative stain images of one-day-old MscS-NCMN particles after SEC separation (White scale bar represents 50 nm).

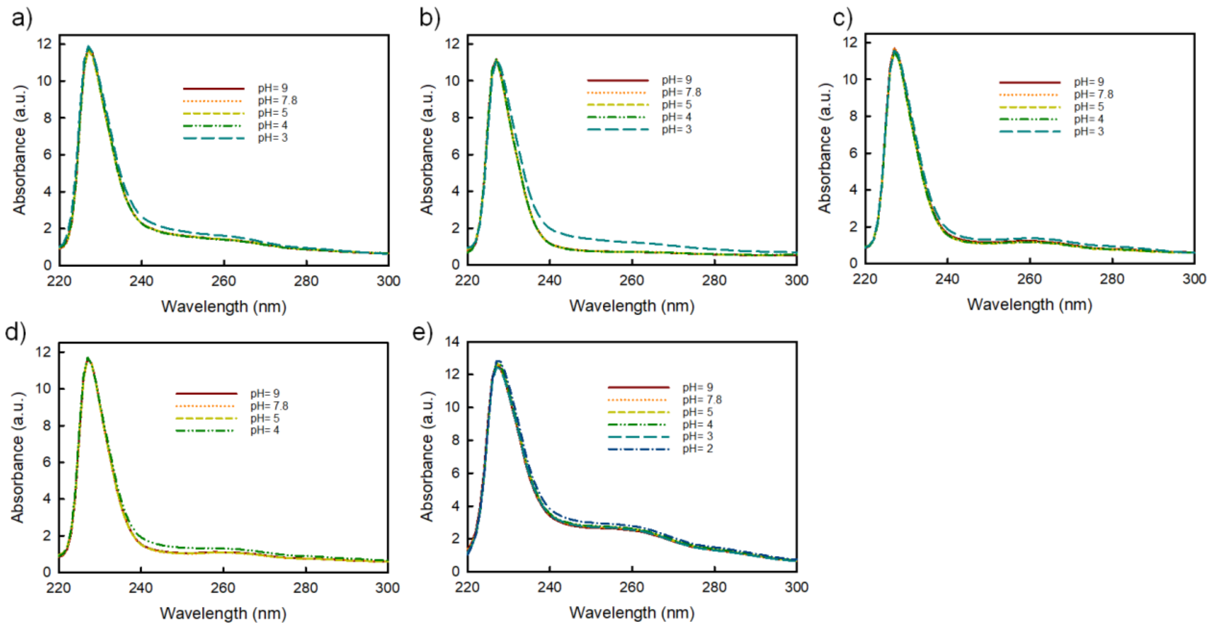

**Figure S6.** Dependency of pH susceptibility of AcrB-NCMN particles on the chemical nature of (a) NCMNP13-50, (b) NCMNP21-20, (c) NCMNP21b-5, (d) NCMNP21b-20 and (e) NCMNP21b-30.

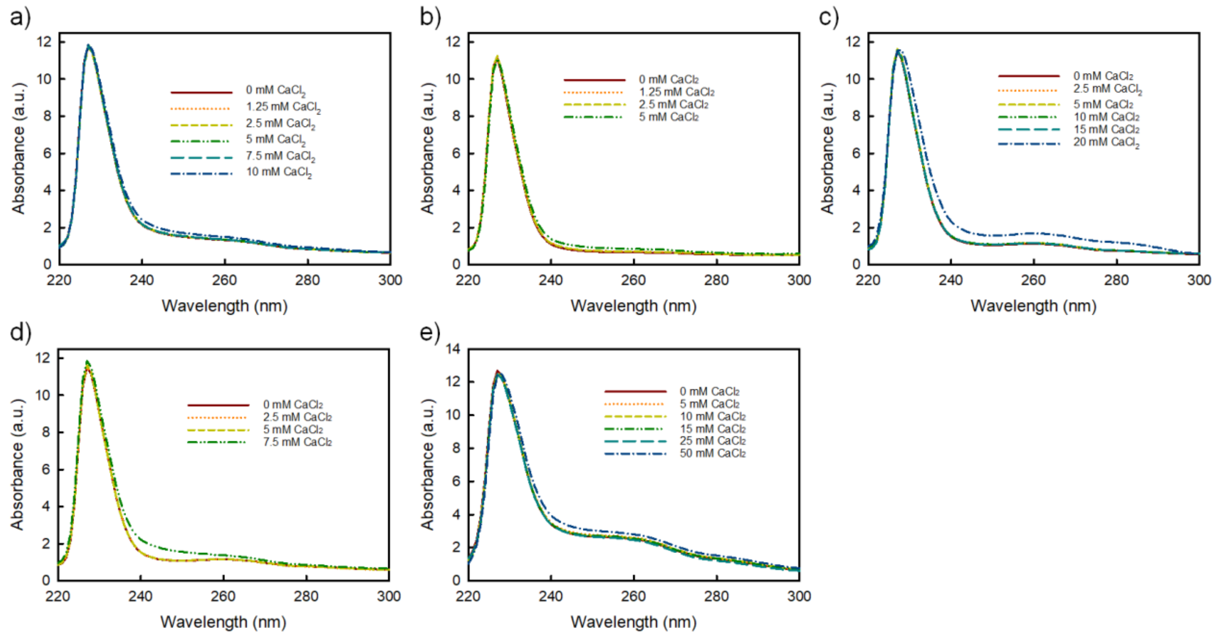

**Figure S7.** Dependency of $Ca^{2+}$ susceptibility of AcrB-NCMN particles on the chemical nature of (a) NCMNP13-50, (b) NCMNP21-20, (c) NCMNP21b-5, (d) NCMNP21b-20 and (e) NCMNP21b-30.

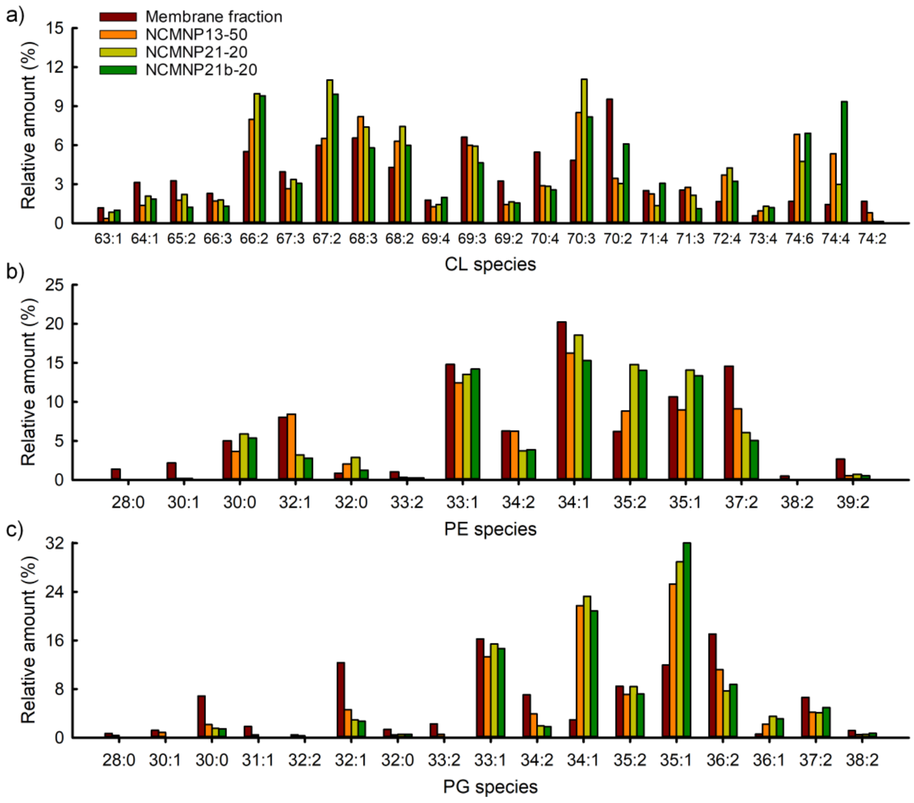

**Figure S8.** Comparison of individual lipid profiles. (a) CL lipid class. (b) PE lipid class. (c) PG lipid class. For each CL, PE and PG lipid class, long alkyl chain lipids with a high degree of unsaturation are enriched in NCMN particles compared with the natural abundance of those lipids on the cell membrane.

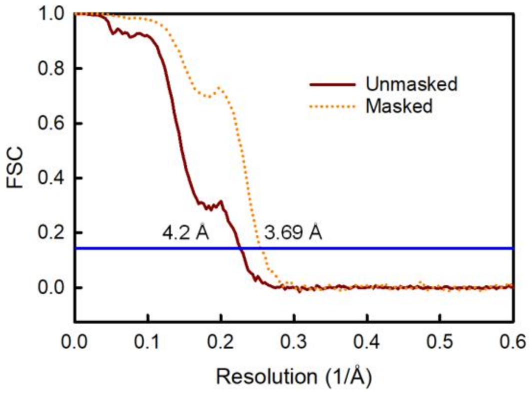

**Figure S9.** The Fourier shell correlation (FSC) resolution graph for 3D reconstruction of AcrB-NCMNP21b-20 using the gold-standard with FSC = 0.143.

#### II. Additional tables

**Table S1.** Cryo-EM data collection and processing

| | **AcrB-NCMNP21b-20** |
|---|---|
| Microscope | TFS Titan Krios |
| Voltage (kV) | 300 |
| Detector | Gatan K3 |
| Nominal magnification | 105,000 × |
| Electron exposure ($e^{-}$ $Å^{-2}$) | 60.22 |
| Defocus range (µm) | 0.4 – 2.9 |
| Pixel size ($Å^{2}$ per pixel) | 0.8256 |
| Dose rate ($e^{-}$/s/pixel) | 44.18 |
| Exposure time (s) | 0.299 |
| Movies stacks (no.) | 3,375 |
| Box size (pixels) | 360 |
| Final particle images (no.) | 86,011 |
| Symmetry imposed | C1 |
| Map resolution | 3.69 Å |
| FSC threshold | 0.143 |
| EMDB ID | EMD-25400 |

#### III. Determination of the grafting levels

As calculated,[11] the average numbers of styrene and carboxylic acid groups of SMA2000 are 18.6 and 16.2, respectively. Therefore, the total aromatic and aliphatic protons of SMA2000 are 98 and 81, respectively. Only the hydrophilic units were altered during our sidechain modification, while the styrene units remained intact. Accordingly, the number of aromatic protons could be used as a reference for calculating the grafting percentage of NCMN polymers. When the broad peak at 5.7 – 8.5 ppm of styrene rings was set at 98 protons, the total aliphatic protons of NCMN polymers were obtained. Based on the difference in aliphatic protons of NCMN polymers and SMA2000, the grafting degree (A%) was determined following the below equations:

- For NCMNP13-x:

$$A\% = \frac{\sum Aliphatic\ proton - 81}{16.2\ (I_{Hb'} + 6I_{Hc'})} x\ 100 = \frac{134.78 - 81}{16.2\ (1+6)}\ x\ 100 = 47.4\ \%$$

- For NCMNP21-x:

$$A\% = \frac{\sum Aliphatic\ proton - 81}{16.2\ (2I_{Hm'} + 2I_{Hn'} + 6I_{Ho'})} x\ 100 = \frac{109.61 - 81}{16.2\ (2+2+6)}\ x\ 100 = 17.7\ \%$$

- For NCMNP21b-x:
    - NCMNP21b-5

$$A\% = \frac{\sum Aliphatic\ proton - 81}{16.2\ (2I_{Hm''} + 2I_{Hn''} + 6I_{Ho''} + 2I_{Hp''} + 2I_{Hq''} + 2I_{Hr''})} x\ 100 = \frac{92.4 - 81}{16.2\ (2+2+6+2+2+2)}\ x\ 100 = 4.4\ \%$$

- NCMNP21b-20

$$A\% = \frac{\sum Aliphatic\ proton - 81}{16.2\ (2I_{Hm''} + 2I_{Hn''} + 6I_{Ho''} + 2I_{Hp''} + 2I_{Hq''} + 2I_{Hr''})} x\ 100 = \frac{127.77 - 81}{16.2\ (2+2+6+2+2+2)}\ x\ 100 = 18.0\ \%$$

- NCMNP21b-30

$$A\% = \frac{\sum Aliphatic\ proton - 81}{16.2\ (2I_{Hm}'' + 2I_{Hn}'' + 6I_{Ho}'' + 2I_{Hp}'' + 2I_{Hq}'' + 2I_{Hr}'')}\ x\ 100 = \frac{152.28 - 81}{16.2\ (2+2+6+2+2+2)}\ x\ 100 = 27.5\ \%$$
